# CaMKII activation triggers persistent formation and segregation of postsynaptic liquid phase

**DOI:** 10.1101/2020.11.25.397091

**Authors:** Tomohisa Hosokawa, Pin-Wu Liu, Qixu Cai, Joana S. Ferreira, Florian Levet, Corey Butler, Jean-Baptiste Sibarita, Daniel Choquet, Laurent Groc, Eric Hosy, Mingjie Zhang, Yasunori Hayashi

## Abstract

Transient information input to brain leads to persistent changes in synaptic circuit, thereby forming memory engrams. Synapse undergoes coordinated functional and structural changes during this process but how such changes are achieved by its component molecules still largely remain enigmatic. We found that activated CaMKII, the central player of synaptic plasticity, undergoes liquid-liquid phase separation (LLPS) with NMDAR subunit GluN2B. Due to CaMKII autophosphorylation, the condensate stably persists even after Ca^2+^ is removed. The selective binding of activated CaMKII with GluN2B co-segregates AMPAR/neuroligin (NLGN) into a phase-in-phase assembly. Because postsynaptic NLGN clusters presynaptic neurexin and other active zone proteins thereby increasing the release probability of synaptic vesicles, this ensures efficient synaptic transmission. In this way, Ca^2+^-induced and persistent formation of LLPS by CaMKII serves as molecular basis of memory by functioning as an activity-dependent crosslinker for postsynaptic proteins and segregating trans-synaptic nanocolumns.

## Introduction

Within a central excitatory synapse, various molecules are segregated into functional nanodomains to accomplish the intricate regulation of synaptic transmission and plasticity. Within the presynaptic compartment, the readily releasable pool of vesicles is concentrated at specialized nanodomains referred as active zones. On the postsynaptic membrane, different classes of glutamate receptor form discrete nanodomains ^1–4^. These pre- and postsynaptic nanodomains are matched with each other, thereby forming trans-synaptic nanocolumns or nanomodules that ensures an efficient transmission between pre- and postsynaptic structures ^1,2,5,6^.

However, how such nanodomains are formed and regulated by neuronal activity, in the absence of any demarcating membranous structures, has not been fully elucidated. Recently, liquid-liquid phase separation (LLPS) of biological macromolecules was found to play a critical role in regulating the assembly and segregation of molecules within various intracellular structures ^7, 8^. In this regard, CaMKII, a highly abundant protein kinase in the postsynaptic density (PSD), has ideal features to undergo LLPS ^7, 8^. Ca^2+^/calmodulin binding to CaMKII opens up a binding pocket called the T-site, which is occupied by the autoinhibitory domain encompassing threonine (T) 286 in inactive kinase, and forms a stable complex with various synaptic proteins at μM affinity, such as the carboxyl tail of NMDA-type glutamate receptor (NMDAR) subunit GluN2B (Fig. S1) and RacGEF protein Tiaml^9, 10^. Once bound, it persists even when cellular Ca^2+^ concentration decreases ^9, 10^. Finally, the dodecameric structure of CaMKII ^11^ allows multivalent interactions.

Given this, we explored whether CaMKII has an ability to undergo LLPS with PSD proteins and, if it does, how it can affect synaptic protein distribution and function. We found that Ca^2+^ activation of CaMKII results in persistent LLPS with PSD proteins in a manner requiring T286 autophosphorylation. CaMKII then segregates two subtypes of glutamate receptor, AMPAR and NMDAR, through the formation of phase-in-phase, which was recapitulated in neurons as revealed by super-resolution microscopy. Neuroligin-1 (NLGN1), a neuronal adhesion molecule, which clusters presynaptic neurexin and other active zone proteins, segregates together with AMPAR. Through these mechanisms, activated CaMKII can undergo persistent LLPS in PSD and establishes AMPAR nanodomain beneath active transmitter release site, thereby conducting a novel mechanism of activity-dependent and persistent synaptic plasticity.

## Results

### CaMKII undergoes LLPS with GluN2B carboxyl tail

In order to test the idea that CaMKII can undergo LLPS with its T-site binding partner, we combined purified CaMKII with carboxyl tail of GluN2B, a prototypical T-site binding protein (residue 1226-1482, GluN2Bc). GluN2Bc was fused with dimeric near-infrared fluorescent protein eqFP670 to label and to mimic the subunit stoichiometry of GluN2B subunit in the endogenous NMDAR complex. We used a low speed centrifugation assay to assess the macromolecular complex formation ^12–14^. Cytoplasmic concentration of CaMKII in the synapse is estimated to be 20-80 μM as a monomer ^15^. Here, we used 10 μM of CaMKII as it was a practical limit of the preparation. Generally, proteins more readily form condensates at higher concentration. Therefore, we are towards the more conservative side in making this conclusion. On the other hand, GluN2B is a membrane protein and it is difficult to define its concentration/density. Also, the association with the membrane limits its diffusion and stability, which can effectively increase the valency of the interaction. Therefore, we tentatively used GluN2B in the same concentration with CaMKII. When CaMKII, GluN2Bc, and calmodulin were mixed in the absence of Ca^2+^, the proteins stayed in the supernatant (Fig. 1A, B). However, upon addition of Ca^2+^, the majority of CaMKII moved to the pellet with GluN2Bc, indicating that Ca^2+^ stimulation of CaMKII induces the formation of a macromolecular complex with GluN2Bc. Differential interference contrast (DIC) and fluorescent microscopy revealed no condensate in the absence of Ca^2+^ (Fig. 1C). However, the addition of Ca^2+^ induced formation of protein condensates containing CaMKII and GluN2Bc, consistent with the sedimentation assay ^12, 14^. Upon point photobleaching within a single condensate, both CaMKII and GluN2Bc fluorescence recovered after photobleaching (Fig. 1D and S2). Once formed, the condensates were stable, and we could observe two droplets fusing together to form a larger droplet (Fig. 1E). These observations indicate that the condensate retained liquid-like properties. GluN2Bc without CaMKII or CaMKII with eqFP670 fusion tag only did not pellet or form condensates indicating that both CaMKII and GluN2Bc are required (Fig. S3A-D). The carboxyl tails of AMPA receptor subunits GluA1 and GluA2 did not form condensates with CaMKII (Fig. S3E). Together, our results indicate that Ca^2+^/calmodulin can trigger formation of protein condensates containing CaMKII and GluN2Bc by a LLPS-mediated mechanism.

**Figure 1.**
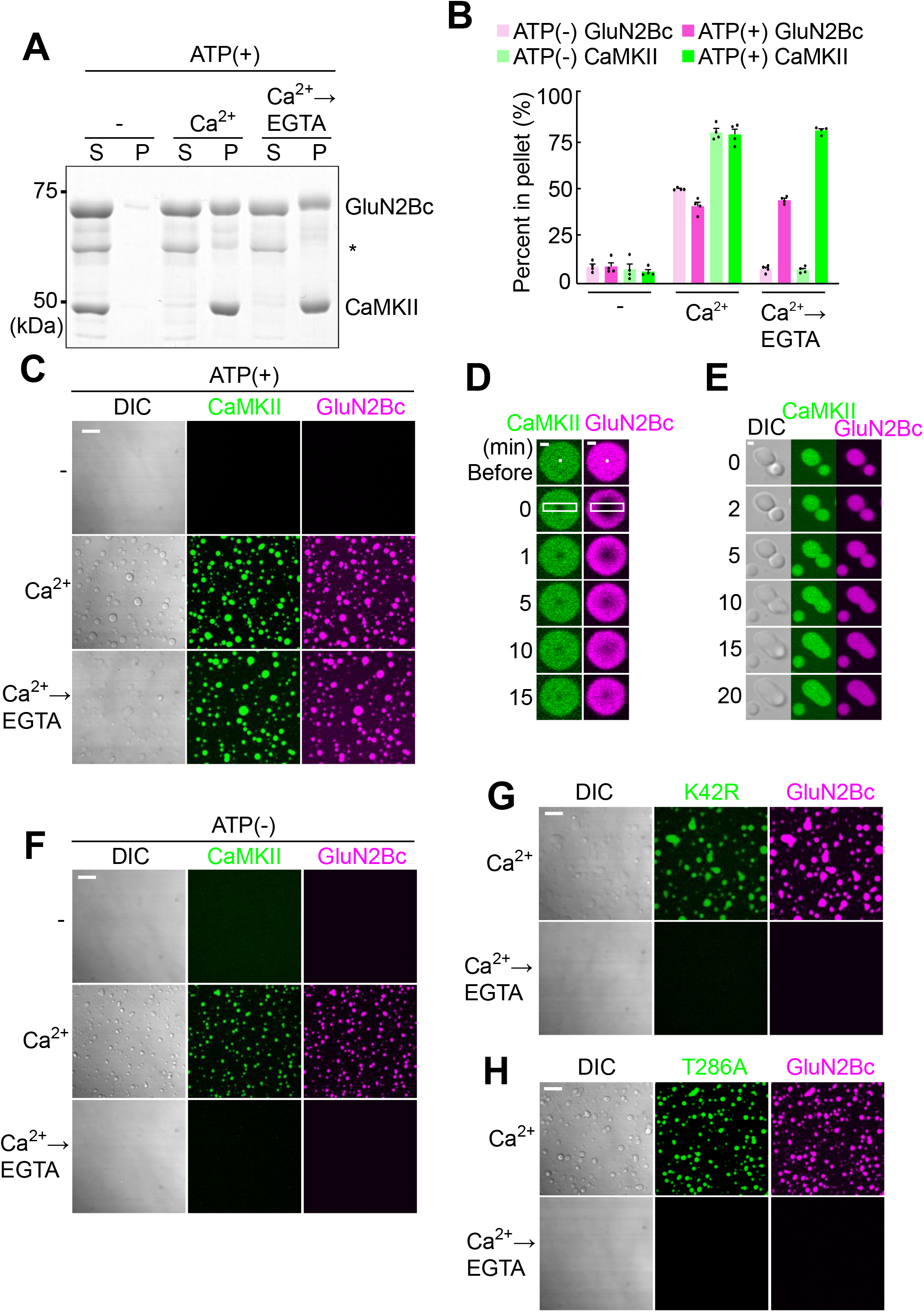
CaMKII and GluN2Bc form LLPS condensates. (A) Low speed sedimentation assay. Ten μM CaMKII and 10 μM GluN2Bc were mixed in the presence of 0.5 mM EGTA, 10 μM calmodulin, 5 mM MgCl_2_ and 2.5 mM ATP (-). Then 2 mM CaCl_2_ (Ca^2+^) was added, followed by 2.5 mM EGTA (Ca^2+^→EGTA). The supernatant (S) and pellet (P) after centrifugation at 10,000 g were subjected to SDS-PAGE and CBB staining. A slight upward shift of GluN2Bc is likely due to phosphorylation by CaMKII. * indicates degradation product of GluN2Bc. Calmodulin is unobservable within the image due to its small size. (B) Quantification of (A) and (S2A) (mean ± SEM, n=4 samples). (C) DIC and confocal microscopic images of the protein mixture as in (A). Only in the presence of Ca^2+^, CaMKII and GluN2Bc formed condensate. Once formed, the condensate persisted even after the addition of EGTA. Scale bar, 10 μm. (D) Fluorescence recovery after photobleaching (FRAP) after photobleaching single point inside of a condensate (indicated by a white dot) of CaMKII-GluN2Bc in the presence of Ca^2+^. Note that they are two separate experiments. Scale bar, 1 μm. See Figure S2 for quantification. (E) A fusion event of condensates. Scale bar, 1 μm. (F) Same experiment as in (C) in the absence of Mg^2+^-ATP. Ca^2+^ triggers the formation of the condensate in the absence of Mg^2+^-ATP. However, if EGTA is added, the condensate was dispersed. Scale bar, 10 μm. (G, H) Same experiment as in (C) using K42R (G) and T286A (H) mutants of CaMKII. Only Ca^2+^ and Ca^2+^→EGTA conditions are shown. In both cases, the condensate was formed in the presence of Ca^2+^ but they did not persist after the addition of EGTA. Combined, the results indicate that T286 phosphorylation is crucial for the persistence of the condensate. Scale bar, 10 μm.

### Autophosphorylation of CaMKII is required for persistence of protein condensate

CaMKII and GluN2Bc protein condensates persisted even after the addition of ethylene glycol tetraacetic acid (EGTA), a Ca^2+^-chelator (Fig. 1A-C). In contrast, in the absence of ATP, condensates could form but dissolved upon addition of EGTA (Fig. 1B, F and S4A). This suggests that the kinase reaction is involved in the persistence of condensates after Ca^2+^ dissipates. Hereafter, all experiments are performed in the presence of ATP unless otherwise indicated. Consistent with the experiment in the absence of ATP, a kinase null CaMKII K42R mutant formed condensates in the presence of Ca^2+^ but failed to persist after the addition of EGTA (Fig. 1G). Mutation of the autophosphorylation site at T286, a site required for the constitutive activation of CaMKII beyond the period of elevated Ca^2+^ concentration, to alanine (T286A) replicated the results of the kinase null mutant (Fig. 1H). These results indicate that the autophosphorylation at T286 is not critical for the initial formation of condensates by Ca^2+^ but is required for the persistent maintenance of the condensates in the absence of Ca^2+^.

We next tested if the multivalent interaction between CaMKII and GluN2Bc is required for the formation of condensates. Consistent with the requirement of multivalency of CaMKII, a catalytically active but monomeric CaMKII mutant 1-314 lacking the association domain failed to form condensates (Fig. S4B). To prevent the specific interaction between CaMKII and GluN2Bc, we turned to a model of binding between CaMKII and GluN2B ^16^. The model shows the interaction between a hydrophobic pocket made by I205 and F98 of CaMKII with L1298 of GluN2B as well as electrostatic interactions between E139 of CaMKII with R1300 of GluN2B. A CaMKII T-site mutant I205K failed to form condensates (Fig. S4C). Also, GluN2Bc mutants which cannot interact with CaMKII, L1298A/R1300Q (LR/AQ) and R1300Q/S1303D (RS/QD) ^9, 17^ failed to form condensates (Fig. S4D, E). These results indicate that multivalent interactions via those hydrophobic and electrostatic interactions between CaMKII T-site and GluN2Bc are required for the formation of condensates.

To obtain temporal information of the formation and the dispersion of condensates, we measured the turbidity of protein mixture (Figure S5)^18^. The turbidity of the CaMKII/GluN2Bc sample increased within 30 sec after the addition of Ca^2+^ and remained after adding EGTA. On the other hand, the turbidity of the T286A/GluN2Bc sample increased similarly to the wildtype CaMKII sample but decreased to the baseline level within 30 sec after EGTA treatment.

### Segregation of AMPAR and NMDAR by LLPS-mediated mechanism of CaMKII

We then added additional components of the PSD to examine if CaMKII can form condensates with other major PSD proteins as well. We tested the carboxyl tail of Stargazin (STGc), an auxiliary subunit of AMPAR critical for determining its synaptic distribution, as a proxy of the AMPA receptor, and PSD-95, which can interact with both GluN2Bc and STGc through PDZ domains ^13, 19, 20^ (Fig. S1). STGc was fused with a tetrameric fluorescent protein DsRed2 to mimic stoichiometry of the endogenous AMPAR complex. When CaMKII, calmodulin, PSD-95, GluN2Bc, and STGc were combined, PSD-95, GluN2Bc, and STGc formed homogenous condensates while CaMKII remained in the diluted phase in the absence of Ca^2+^ (Fig. 2A-C) ^20^. However, when Ca^2+^ was added, CaMKII partitioned into the condensates, which persisted after the addition of EGTA. Intriguingly, we found a segregation of proteins within the condensate. CaMKII and GluN2Bc came to the periphery and surrounded PSD-95 and STGc, which formed a phase-in-phase organization (Fig. 2C). Z-axis reconstruction revealed that CaMKII and GluN2Bc entirely covered PSD-95 and STGc (Fig. 2D). While STGc was exclusively enriched in the inner phase, PSD-95 was partitioned in the peripheral phase as well (Fig. 2E). Conversely, both CaMKII and GluN2Bc were also partly partitioned in the inner phase as well. Again, consistent with liquid-like properties, we observed the condensates fusing with each other (Fig. 2F).

**Figure 2.**
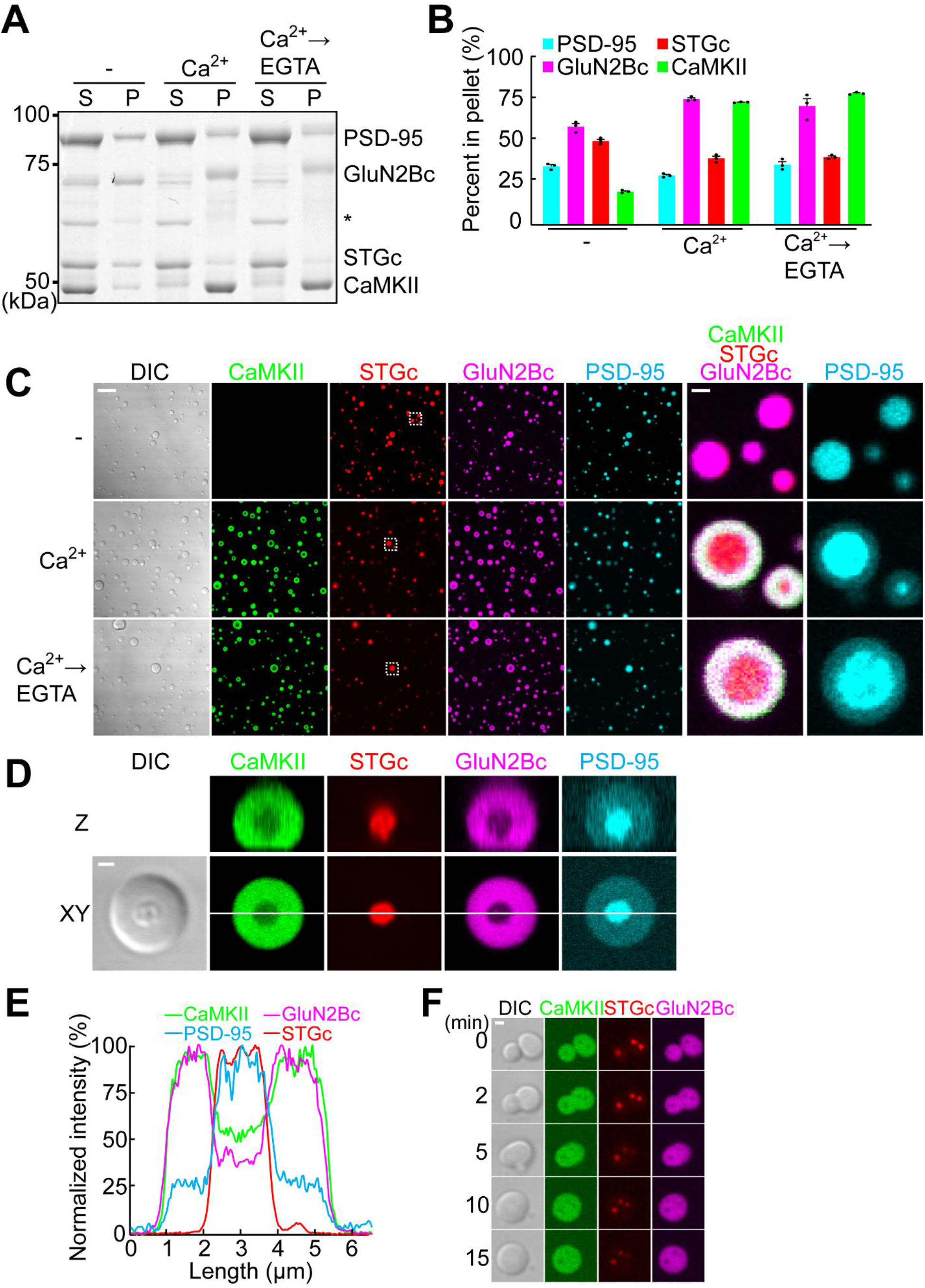
Segregation of AMPAR and NMDAR within protein condensate by active CaMKII. (A) Sedimentation assay of 10 μM PSD-95, 2.5 μM GluN2Bc, 7.5 μM STGc, 10 μM CaMKII, and 10 μM calmodulin in the presence of Mg^2+^-ATP. The upward shift of band and the reduction in the staining of PSD-95, GluN2Bc, and STGc is likely due to phosphorylation by CaMKII. (B) Quantification of (A) (mean ± SEM, n=3 samples). (C) Images of the protein mixture as in (A). Right two columns are high magnification of the dashed rectangle in the STGc channel. Scale bars, 10 μm and 1 μm for low- and high-magnification images. (D) Magnification and Z projection of single condensates. Scale bar, 1 μm. (E) Line scanning of (D) in each color channel. (F) Observation of a condensate fusion event. Scale bar, 1 μm. When stimulated with Ca^2+^, PSD-95/STGc formed phase-in-phase while GluN2Bc/CaMKII formed a surrounding phase. This persisted even after addition of EGTA.

The formation of the phase-in-phase organization requires CaMKII. Without CaMKII, Ca^2+^ failed to induce the phase-in-phase assembly (Fig. S6A). In the presence of CaMKII, the phase-in-phase assembly could be induced in the absence of ATP (Fig. S6B). However, after addition of EGTA, CaMKII moved to the diluted phase and the condensates became homogenous (Fig. S4B). Essentially the same results were obtained by using the CaMKII K42R (Fig. S6C) and T286A mutants (Fig. S7A) in the presence of ATP. These results indicate that neither kinase activity nor T286 phosphorylation is required for the phase-in-phase assembly formation even though GluN2B, Stg, and PSD-95 are all known to be phosphorylated by CaMKII ^9, 19, 20^. However, for the persistent phase-in-phase organization after the decrease in Ca^2+^ concentration, T286 autophosphorylation is crucial. CaMKII I205K and 1-314 mutants did not induce segregation (Fig. S7B and C). Together, these results indicate that the segregation of GluN2Bc and STGc requires multivalent binding at the CaMKII T-site and GluN2Bc, but not the phosphorylation of any of the components. However, the persistent segregation after Ca^2+^ receding requires CaMKII T286 phosphorylation.

### T-site interaction peptide can dissolve the protein condensates

Different synaptic input patterns can induce bidirectional synaptic plasticity. We then wondered if there is any way to reverse the protein condensates. We turned to Camk2n1, a small endogenous CaMKII inhibitor protein which interacts with the T-site of CaMKII and is upregulated during memory processes ^21^. Infusion of the Camk2n1 to protein condensates resulted in collapse of the condensates (Fig. 3A, B. Movies 1 and 2). In the condensates composed of CaMKII/GluN2Bc/PSD-95/STGc, the surrounding CaMKII/GluN2Bc phase collapsed, while the PSD-95/STGc in the inner phase was more resistant, consistent with the fact that PSD-95/STGc by themselves form condensates ^20^ (Fig. 3B). To confirm Camk2n1 disrupts the phase by competing with the T-site, we used CN21, a peptide derived from the minimum T-site binding region of Camk2n1 ^22^. CN21, but not a scrambled peptide, collapsed the condensates formed by CaMKII and GluN2Bc (Fig. 3C). Although CN21 is a CaMKII inhibitor, in this case, it does not affect existing phosphorylation as there is no phosphatase. These results indicate that the LLPS mediated by CaMKII can be reversed by Camk2n1 competition with GluN2Bc.

**Figure 3.**
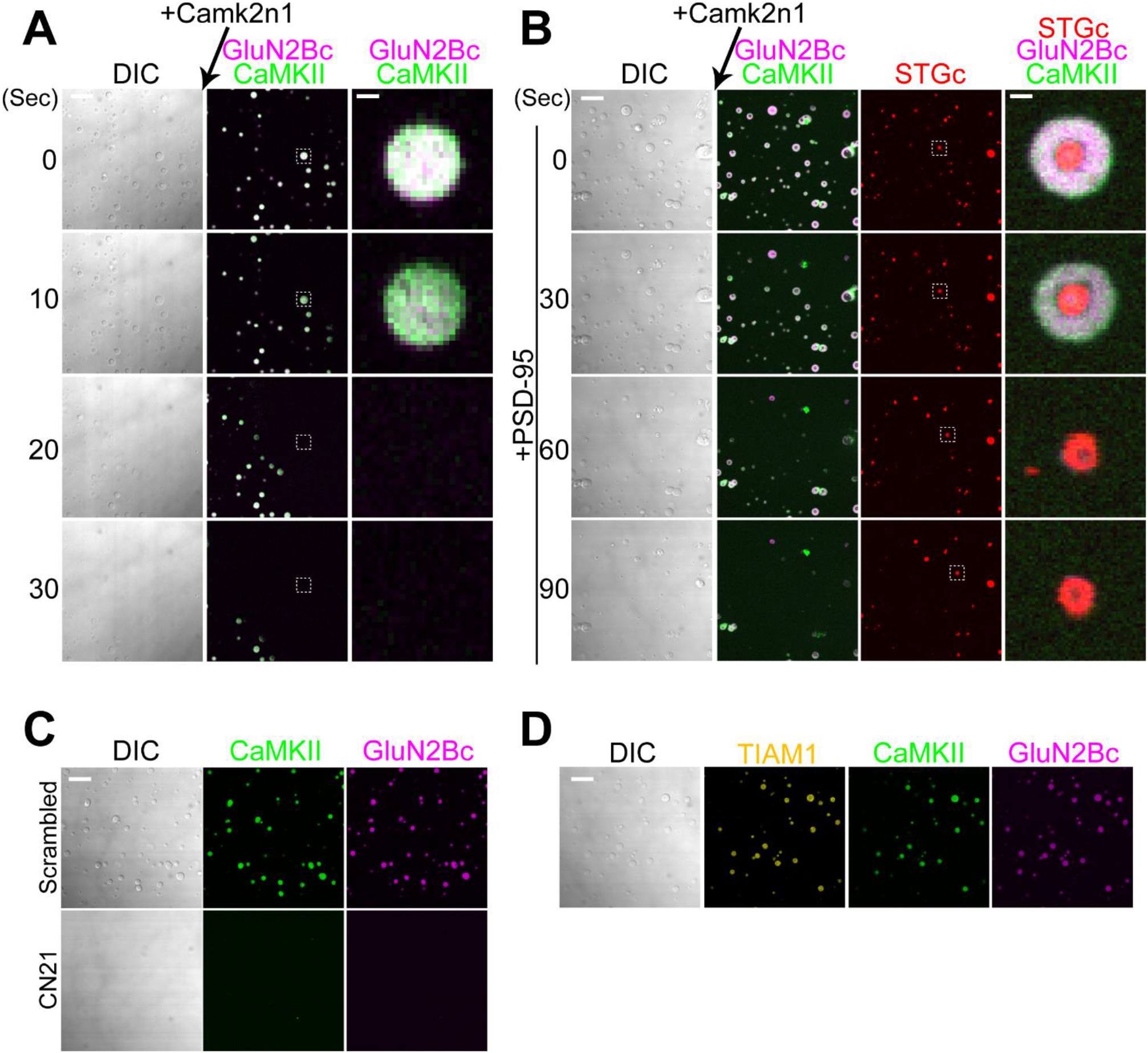
Dispersion of protein condensates by competing T-site interaction. (A) Time-lapse imaging of CaMKII-GluN2Bc condensates (Ca^2+^→EGTA condition) during infusion of 50 μM Camk2n1. Arrow shows the direction of infusion. See Movie 1. Scale bars, 10 μm and 1 μm for low- and high-magnification images. Note a complete dispersal of the condensate. (B) Same experiment as in (A) using the condensates of CaMKII, GluN2Bc, PSD-95 and STGc. Due to the limited number of color channels available, PSD-95 was not imaged. See Movie 2. Note that the phase-in-phase of PSD-95/STGc was resistant to Camk2n1 application, indicating that these two proteins formed condensates by themselves. (C) Effect of 5 μM scrambled and CN21 peptides for CaMKII/GluN2Bc condensates in Ca^2+^→EGTA condition. CN21 replicated the effect of Camk2n1. (D) Effect of 5 μM Tiam1 peptides for CaMKII/GluN2Bc condensates in Ca^2+^→EGTA condition. Tiam1 peptide was taken up by the condensate without much affecting the LLPS. Scale bars,10 μm and 1 μm for low- and high-magnification images.

### Disruption of CaMKII T-site interaction decreases segregation between AMPAR and NMDAR

We then tested if CaMKII plays a role in segregating AMPAR and NMDAR in neurons by using direct stochastic optical reconstruction microscopy (dSTORM) ^3, 4^. We immunolabeled endogenous AMPAR subunit GluA2 and NMDAR subunit GluN1 by using antibodies against their extracellular domains and then analyzed the overlap of synaptic nanodomains between the two receptor subtypes. In control neurons treated with cell-permeable peptide tat-scrambled (SCR), we could observe AMPAR and NMDAR form distinct nanodomains (Fig. 4A). In neurons treated with tat-CN21 (CN21), the overlap was significantly increased as compared to those treated with SCR, consistent with the idea that the segregation of AMPAR and NMDAR is dependent on CaMKII-mediated phase-in-phase assembly formation (Fig. 4B, C). The reason why the proteins did not totally diffuse away by CN21 treatment unlike the LLPS experiment is likely due to the presence other multiple mechanisms that still anchor the receptors at the synapse. We did not find a change in the area of nanodomain, the number of localization and the density of localization in CN21 treated neurons compared with the neurons treated with the SCR (Fig. S8A-C).

**Figure 4.**
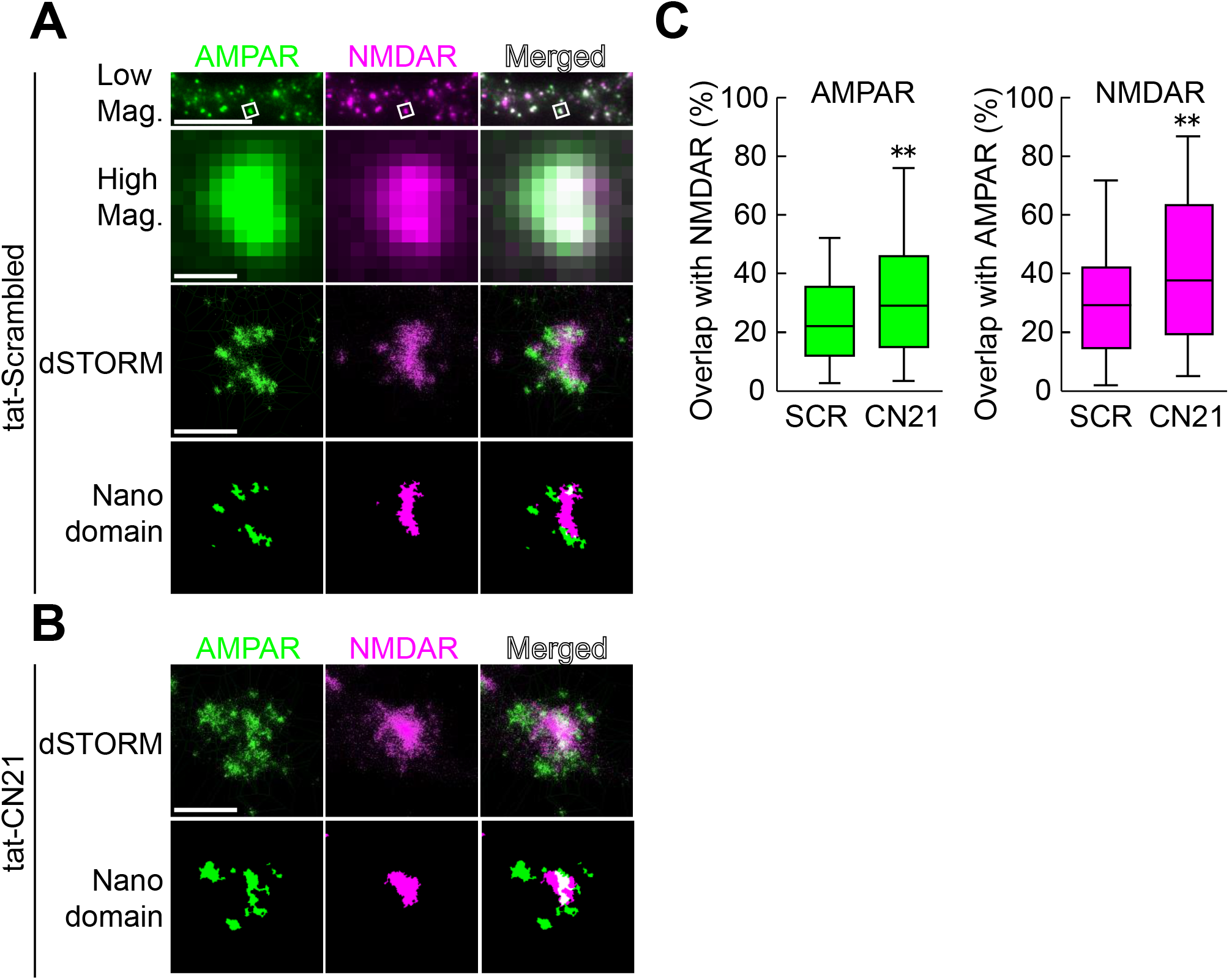
Reduction of synaptic glutamate receptor segregation by competing T-site interaction. (A) From top to bottom. Low magnification epifluorescence image of a dendrite from a hippocampal neuron in dissociate culture treated with 20 μM tat-scrambled peptide for 30 min and immunolabeled with anti-GluA2 (AMPAR, green) and anti-GluN1 (NMDAR, magenta). Scale bar, 10 μm. High magnification image of a single synapse (in white squares in the low magnification image). dSTORM and thresholded images of the same region. Scale bar, 0.5 μm. (B) Images of a synapse treated with 20 μM tat-CN21 for 30 min. (C) Proportion of AMPAR nanodomains overlapping with NMDAR nanodomains (left, p=0.0098) and of NMDAR nanodomains overlapping with AMPAR nanodomains (right, p=0.0019) in tat-scrambled (SCR) or tat-CN21 (CN21) treated neurons. There was significantly more overlap in two receptor nanodomains in neurons treated with tat-CN21 than those treated with tat-scrambled. The data set was obtained from 118 spines (SCR) and 116 spines (CN21), from a total of 10 neurons for each treatment group. All neurons were processed in parallel using the same staining, acquisition and analysis parameters in blind fashion. The statistical significance of the results was assessed by two-sided Mann-Whitney U test. ** indicates p < 0.01.

### Tiam1 behaves as a Ca^2+^-dependent client for CaMKII condensate

We then test the possibility that other synaptic T-site proteins might serve as a client for the CaMKII/GluN2Bc condensate. We previously found that persistent binding between CaMKII and Tiam1, a RacGEF protein, after LTP induction results in a reciprocally-activating kinase-effector complex (RAKEC), which supports persistent Rac activity and the enlargement of dendritic spines ^10^. We therefore tested if fluorescently labeled Tiam1 peptide corresponding to the CaMKII-binding domain (1544-DSHASRMAQLKKQAALSGINGG-1565), can be taken up by the protein condensate (Fig. 3D). As a result, we found that peptide was taken up by CaMKII/GluN2Bc condensates formed by the addition of Ca^2+^. Once taken up, the peptide remained even after Ca^2+^ was chelated. This suggests that the protein condensate formed by CaMKII can serve as a mechanism to trap synaptic T-site binding proteins in an activity dependent fashion.

### NLGN co-segregates together with AMPAR

The trans-synaptic nanocolumn composed of presynaptic active zone and postsynaptic glutamate receptor is refined by neuronal activity, which can enhance the efficiency of synaptic transmission ^2, 5, 23, 24^. We wondered whether CaMKII-mediated segregation of postsynaptic proteins can communicate with the presynaptic terminal. We thus turned to examining the role of neuroligin-1 (NLGN), a neuronal adhesion molecule. NLGN interacts with presynaptic neurexin through its N-terminal extracellular domain, while the intracellular C-terminus interacts with the third PDZ domain (PDZ3) of PSD-95^23, 25–27^ (Fig. S1). We fused the carboxyl tail of NLGN (NLGNc) with dimeric Kusabira Green, a fluorescent protein, and tested if it could form condensates. NLGNc alone (not shown) or together with PSD-95 did not form condensates (Fig. 5A). Only when we added GluN2Bc or GluN2Bc and STGc, the NLGNc participated in condensates (Fig. 5A). The deletion of the PDZ domain binding motif of NLGNc (NLGNc-Δ7) excluded it from the condensates (Fig. S9). These results indicate that NLGNc participates in the PSD-95 condensates as a “client” through its PDZ-binding motif. When we added CaMKII, before addition of Ca^2+^, proteins other than CaMKII formed homogenous condensates (Fig. 5B with unlabeled CaMKII, Fig. 5C with unlabeled PSD-95). Upon stimulation with Ca^2+^, NLGNc moved to the inner phase together with STGc/PSD-95 whereas GluN2Bc and CaMKII segregated to the outer phase (Fig. 5B, C and S10). These results indicate that NLGN is partitioned together with AMPAR and forms a phase distinct from NMDAR, which might serve as a mechanism to position AMPAR beneath the presynaptic active zone. To test if the segregation of NMDAR and NLGN1 in neurons also depends on CaMKII, we treated the neurons in dissociated culture with tat-CN21 or tat-scrambled peptides, surface-labeled and observed them by dSTORM (Fig. S11). In neurons treated with tat-Scrambled peptide, NMDAR and NLGN1 were segregated from each other. In contrast, in neurons treated with tat-CN21, the segregation between them became significantly smaller.

**Figure 5.**
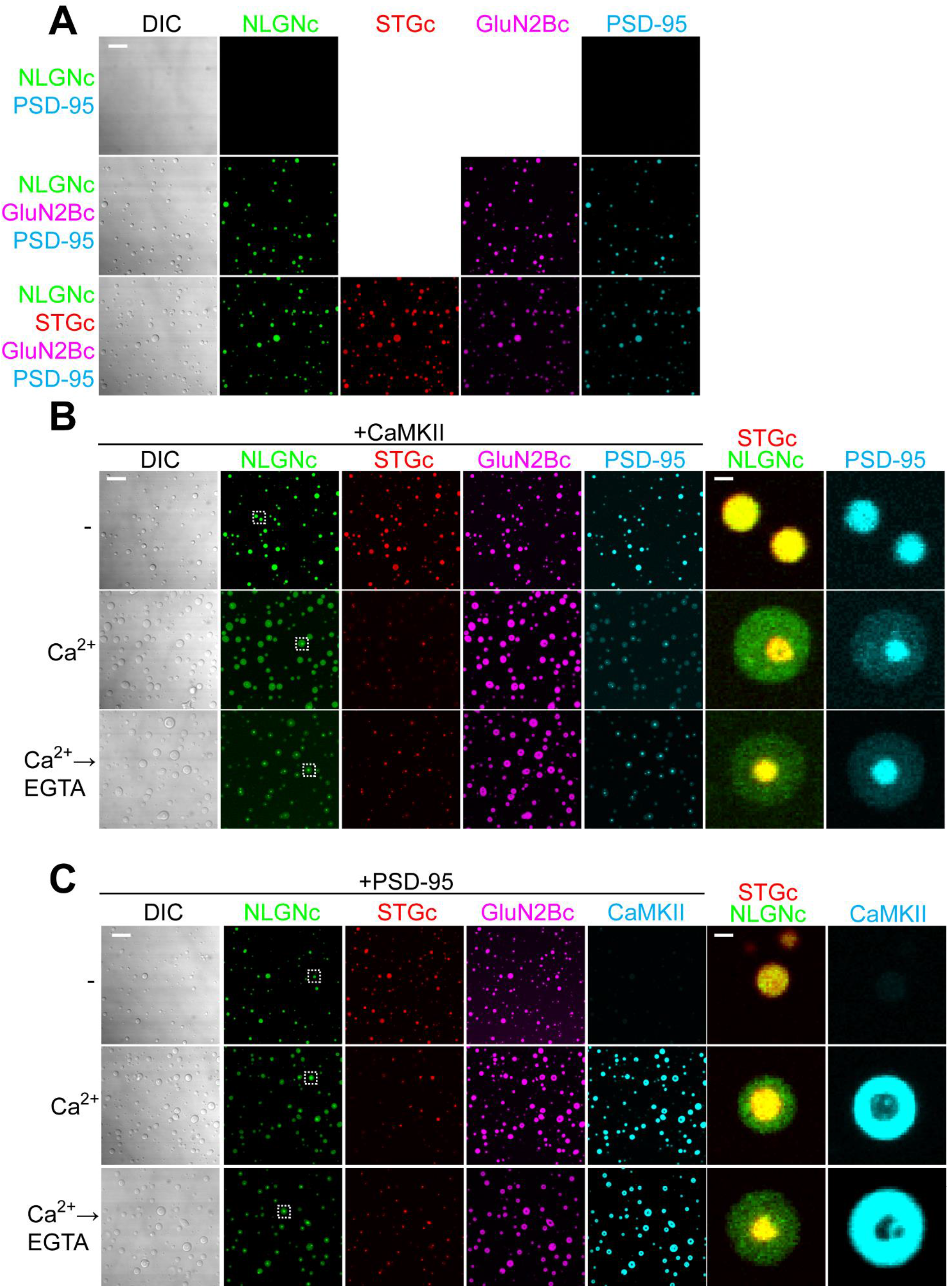
Neuroligin-1 segregates into the STGc/PSD-95 phase by CaMKII. (A) Images of indicated protein mixtures. Ten μM PSD-95 was mixed with 10 μM NLGNc (upper) and an additional 10 μM GluN2Bc (middle) and 10 μM STGc (lower). No Ca^2+^ was added. NLGNc plus PSD-95 absence of GluN2Bc did not form condensate. This indicates that NLGNc participates in the condensates formed by PSD-95 and GluN2Bc as a “client”. (B) Condensates of 10 μM PSD-95, 2.5 μM GluN2Bc, 7.5 μM STGc, 1 μM NLGNc and unlabeled CaMKII in -, Ca^2+^, and Ca^2+^→EGTA conditions. Right column shows higher magnification image of condensates, indicated by the dashed rectangle in NLGNc channel. Due to the limited number of color channels available, we used unlabeled CaMKII and labeled PSD-95. (C) Same experiment as in (B) but using labeled CaMKII and unlabeled PSD-95. Scale bars, 10 μm and 1 μm for low- and high-magnification images. NLGNc/STGc segregate from CaMKII/GluN2Bc to form phase-in-phase while CaMKII and GluN2Bc in surrounding phase in the presence of Ca^2+^. Once formed, the phase-in-phase organization remained even after EGTA was added.

## Discussion

In this study, we revealed that CaMKII can undergo LLPS with major PSD proteins, most notably GluN2B, through its multivalent interaction conferred by its dodecameric structure. This view is consistent with several properties of synaptic CaMKII such as constant exchange at rest as revealed by FRAP analysis ^28^, and rapid translocation to the synapse upon LTP induction in a manner requiring the interaction of CaMKII T-site with GluN2B carboxyl tail ^17, 29, 30^.

The initial formation of protein condensates was triggered by Ca^2+^ and was independent of kinase activity. Once formed, the condensate persisted even after the decrease in Ca^2+^ concentration. For this persistence, CaMKII T286 autophosphorylation is required, which maintains CaMKII in an open conformation and exposes the T-site ^31^, thereby allowing the binding of GluN2B. In its absence, the autoinhibitory domain docks at the T-site ^11^ and competes out the binding with GluN2B. We speculate this is the reason why T286 phosphorylation is required for the persistence of protein condensates.

Within condensates, CaMKII segregated AMPAR and NMDAR into different compartments. Super-resolution imaging of the native AMPAR and NMDAR provided *in vivo* evidence that CaMKII segregates these two subtypes of glutamate receptors into different nanodomains (Fig. 6A). AMPAR partitions together with NLGN and can form a link with the presynaptic active zone. This mechanism may enrich AMPAR beneath the transmitter release site. AMPAR has a low affinity to glutamate compared with NMDAR and is normally not saturated with glutamate at the synaptic cleft ^32–34^. Indeed, super-resolution imaging studies revealed the alignment of pre- and postsynaptic markers, termed synaptic nanomodules or nanocolumns, is refined as a result of neuronal activation ^5, 6^. The segregation of AMPARs under the transmitter release site can increase the efficacy of synaptic transmission (Fig. 6B) ^24^. Furthermore, cluster formation of NLGN induces clustering of presynaptic neurexin, which then recruits additional axonal vesicular release machinery and eventually forms active zone ^35^. Therefore, the postsynaptic clustering of NLGN may serve as a retrograde mechanism to increase presynaptic release probability (Fig. 6B) ^27^. These combined, postsynaptic activation of CaMKII and resultant formation of LLPS can serve as a novel modulatory mechanism of synaptic transmission. Consistent with this idea, the activation of postsynaptic NMDAR accumulates more the active zone proteins over postsynaptic PSD-95 cluster, thereby forms a trans-synaptically matched nanocolumn of release machinery and receptor complex at the synapse ^5^.

**Figure 6.**
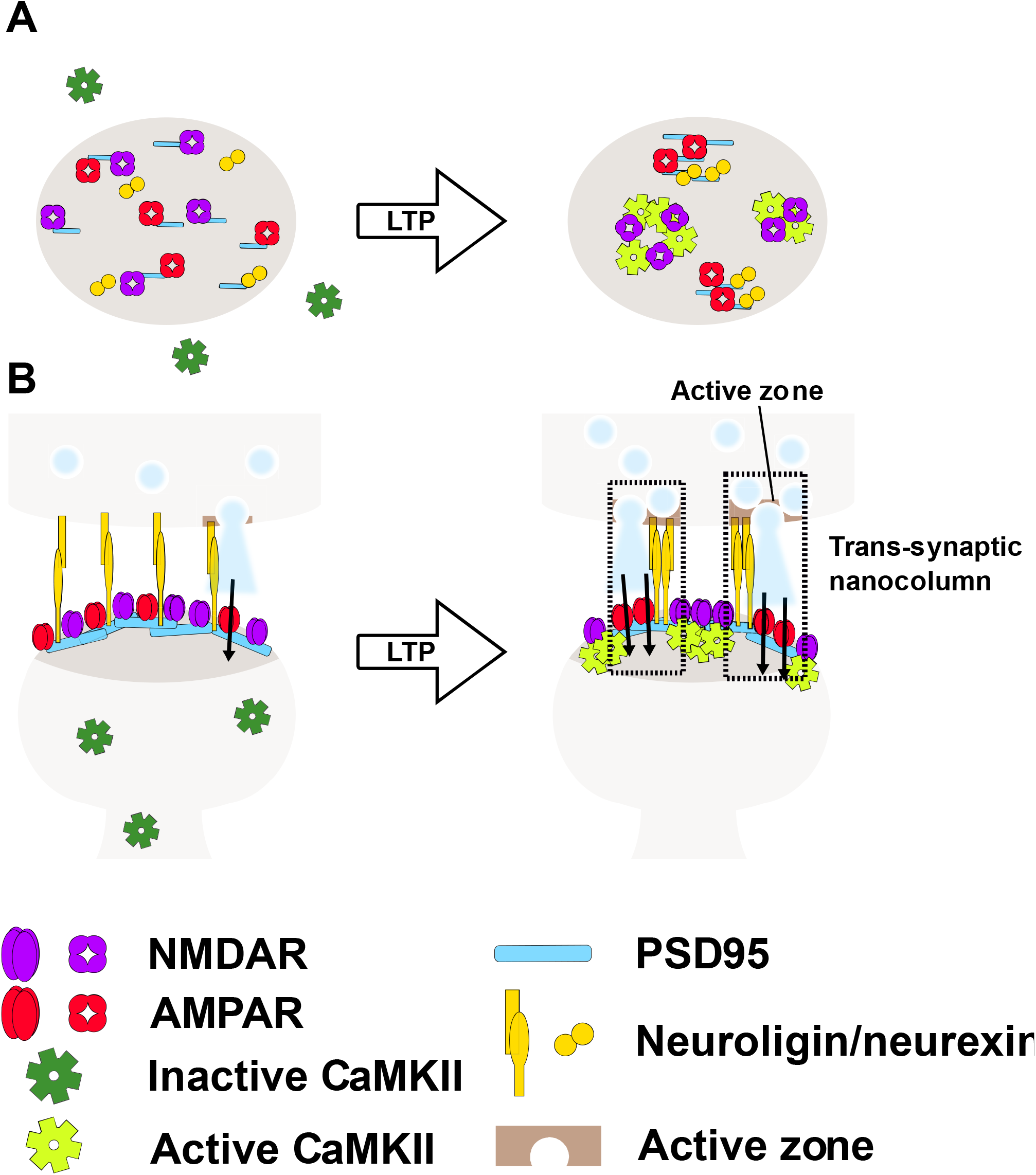
The role of CaMKII for modulation of subsynaptic segregation of glutamate receptors. (A) A top-down view on the postsynaptic membrane. Basally, AMPAR and NMDAR are mixed (left). By the activation of CaMKII, AMPAR is segregated from NMDAR (right). (B) Basally, some of AMPAR are silent because they do not receive sufficient glutamate to open the channel (left). CaMKII-mediated segregation of AMPAR and neuroligin aligns the presynaptic release site and postsynaptic AMPAR nanodomains to increase the efficacy of synaptic transmission (right).

GluN2B is a minor component of PSD proteins ^36^ and the CaMKII T-site can interact with other proteins such as Tiam1, GJD2/connexin 36, LRRC7/densin-180, Camk2n1, and L-type Ca^2+^ channel. Therefore, it is possible that CaMKII forms condensates with these proteins as well, even though GluN2B would be the most important partner for CaMKII ^17^. Through this mechanism, CaMKII can serve as a postsynaptic Ca^2+^-dependent hub, which underlies the activity-dependent transport and crosslinking of multiple postsynaptic client proteins observed during LTP via the LLPS-mediated mechanism ^30, 37^. This reasonably explains the dodecameric structure and abundance of CaMKII.

In conclusion, we proposed a novel mechanism for synaptic plasticity mediated by liquid-liquid phase separation initiated by CaMKII. In the future, the relative contribution of this versus other proposed mechanisms of synaptic plasticity mediated by CaMKII such as AMPAR phosphorylation, insertion and translocation is to be determined ^38–40^.

---

**Supplementary Figure 1.**
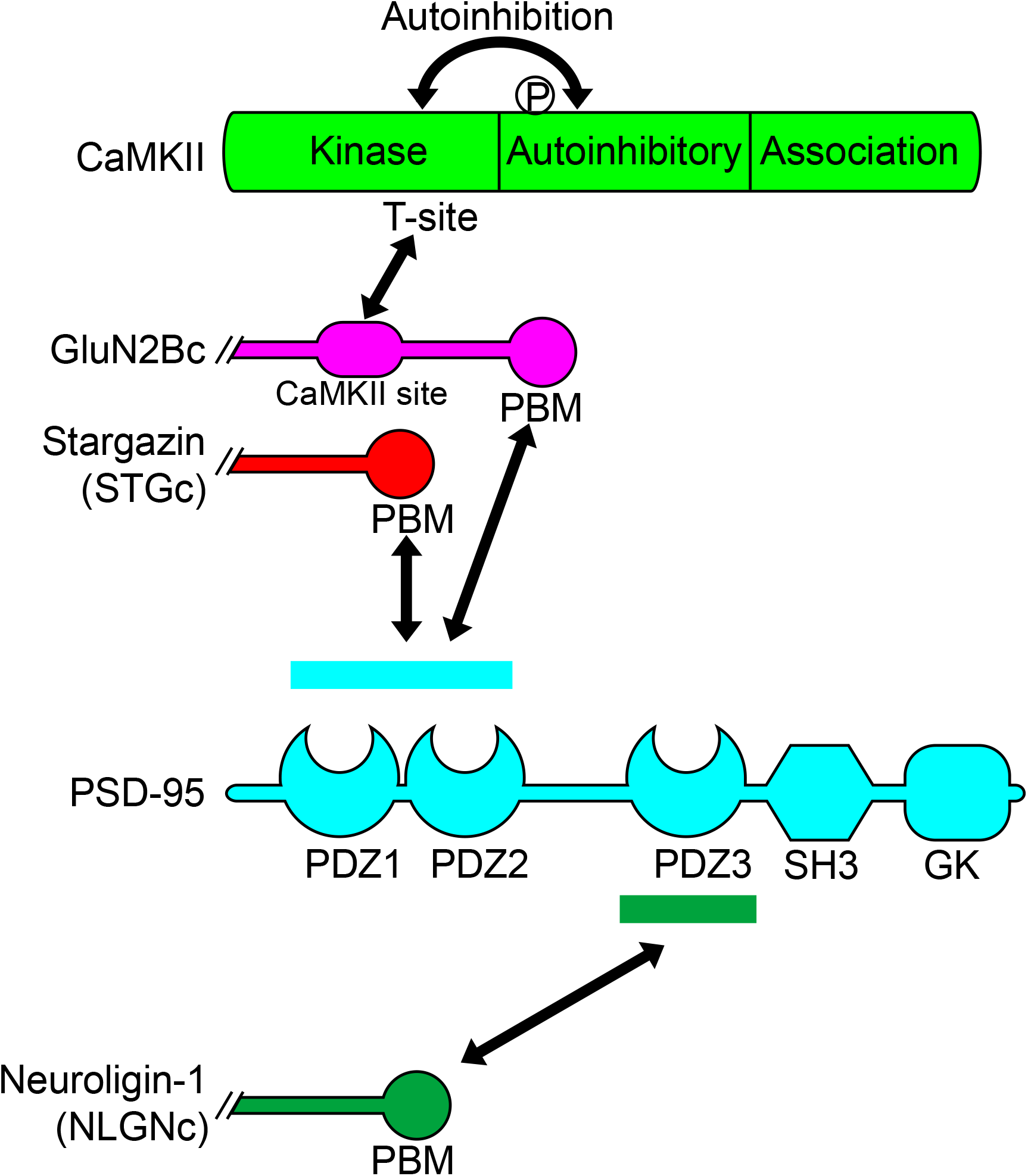
Interaction of proteins used in this study. P denotes T286 autophosphorylation site that renders CaMKII constitutively active. PBM: PDZ-binding motif.

**Supplementary Figure 2.**
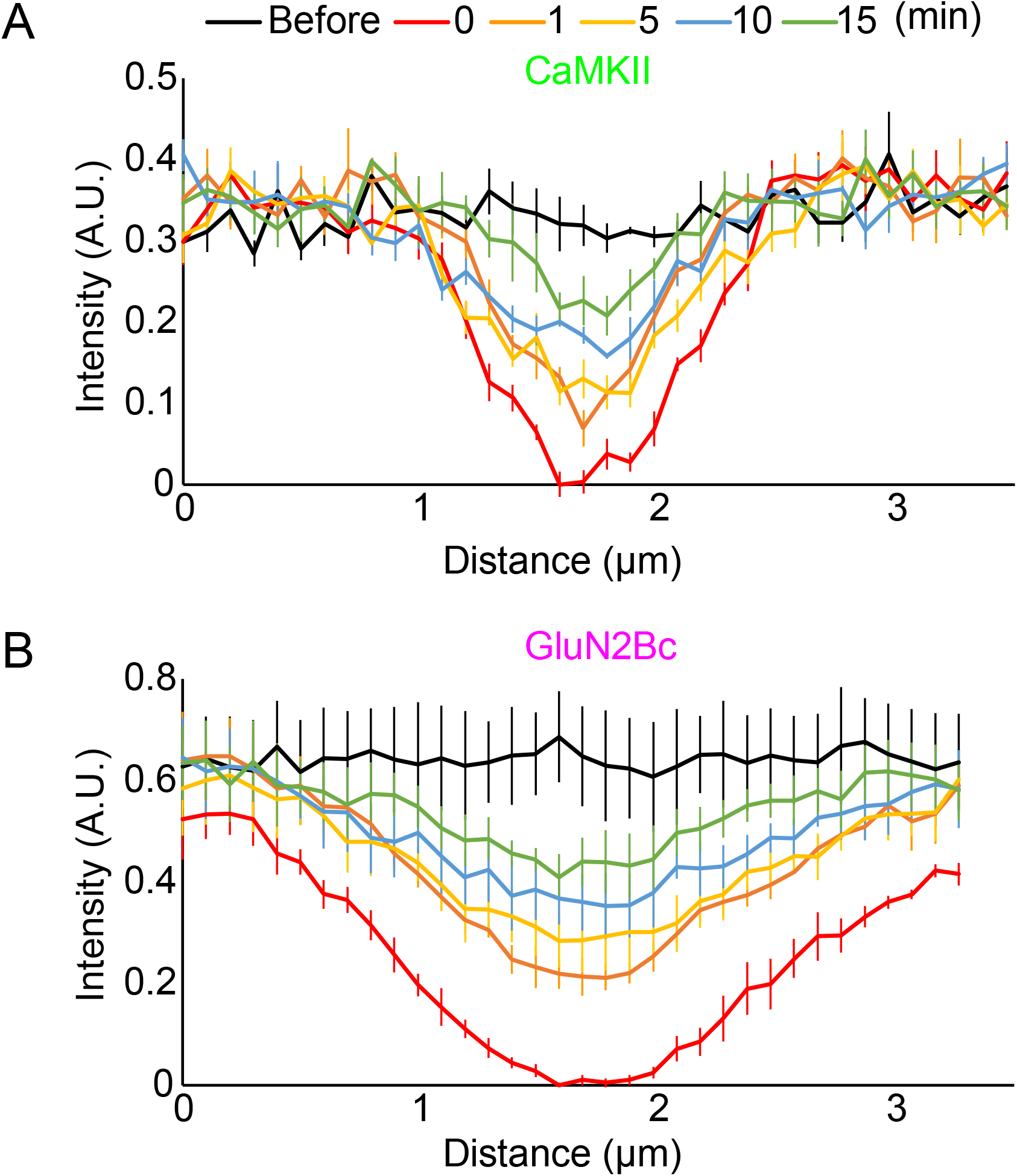
Quantification of fluorescent intensity in Figure 1D. Graphs show the average fluorescent intensity of CaMKII (A) and GluN2Bc (B) across white horizontal line in Fig. 1D from 4 condensates. A.U. arbitrary unit.

**Supplementary Figure 3.**
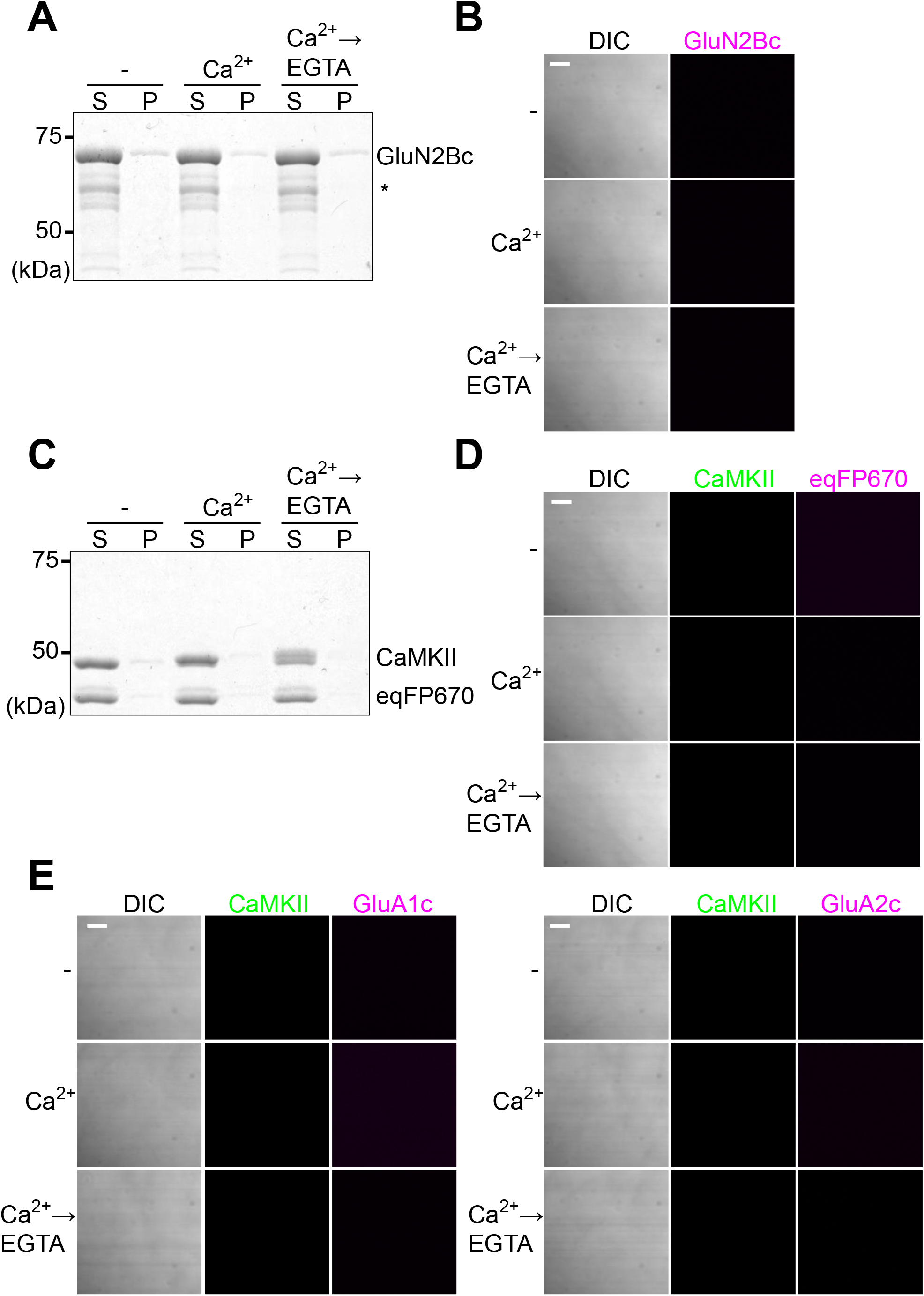
Requirement of CaMKII and GluN2Bc but not AMPAR carboxyl tails for the formation of protein condensates. (A) Sedimentation assay with 10 μM GluN2Bc in the presence of Mg^2+^-ATP. (B) Images of the same protein solution as (A). These results indicate that GluN2Bc alone is not sufficient to undergo LLPS. (C) Sedimentation assay with 10 μM CaMKII and 10 μM eqFP670-SpyCatcher in the presence of Mg^2+^-ATP. eqFP670-SpyCatcher is a fluorescent protein used for labeling GluN2Bc in the rest of the study. This indicates that CaMKII alone is not sufficient to undergo LLPS. (D) Images of the same protein mixture as (C). (E) Images of 10 μM CaMKII with carboxy tails of GluA1 (left) and GluA2 (right) fused with E2-Crimson in the presence of Mg^2+^-ATP. The carboxyl tails of AMPAR did not undergo LLPS with CaMKII. Scale bars, 10 μm.

**Supplementary Figure 4.**
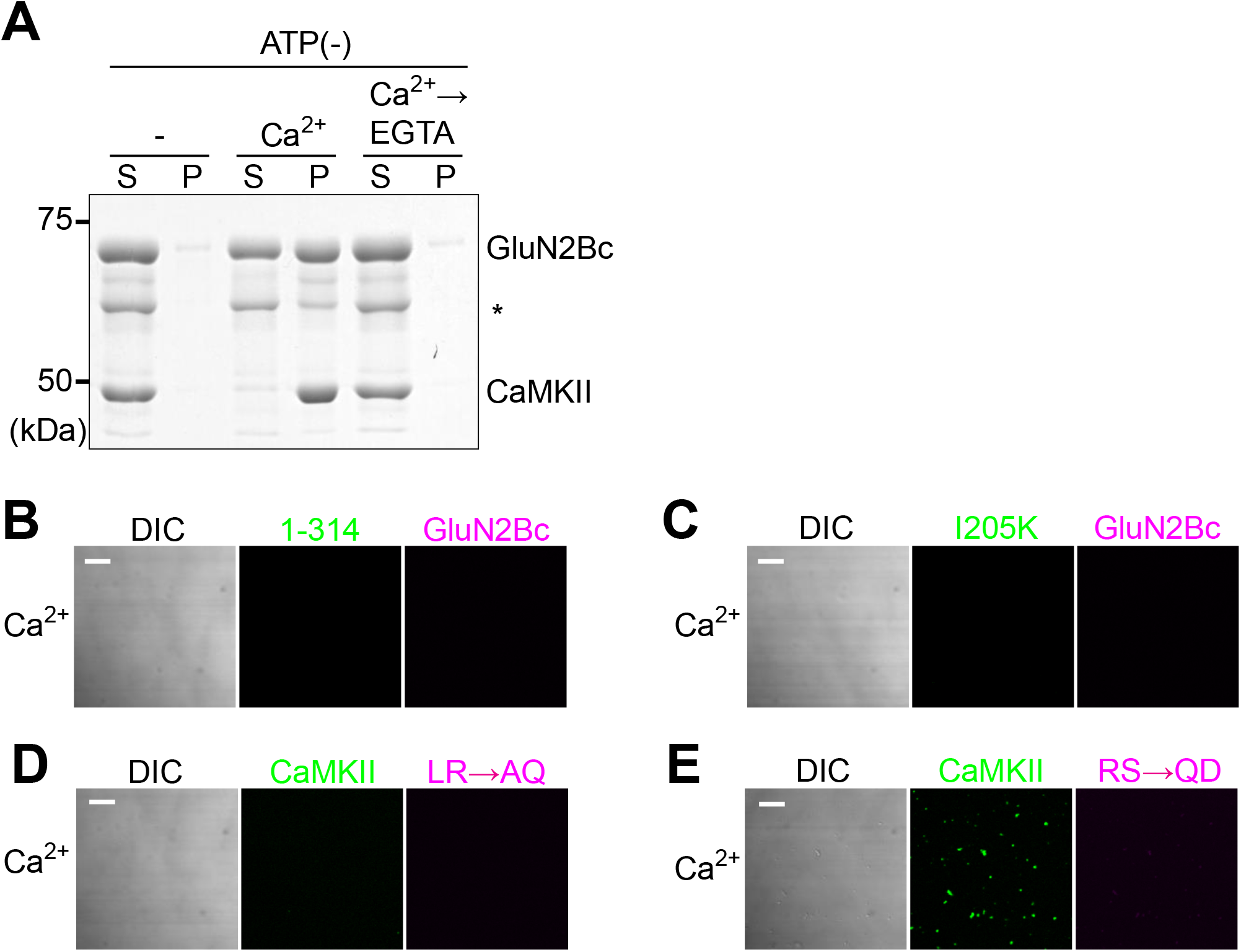
Multivalent interaction and autophosphorylation is required for the formation and persistence of condensates. (A) Sedimentation assay similar to Figure 1A but carried out in the absence of Mg^2+^-ATP. (B, C) Images of CaMKII monomeric 1-314 mutant (B) and T-site mutant I205K (C) each at 10 μM and 10 μM GluN2Bc. Only Ca^2+^ condition is shown. (D, E) Images of 10 μM CaMKII and GluN2Bc CaMKII-binding site mutants L1298A/R1300Q (LR/AQ) (D) and R1300Q/S1303D (RS/QD) (E) each at 10 μM. Only Ca^2+^ condition is shown. Scale bars, 10 μm. These results indicate multivalent interaction between CaMKII T-site and GluN2Bc is required for LLPS.

**Supplementary Figure 5.**
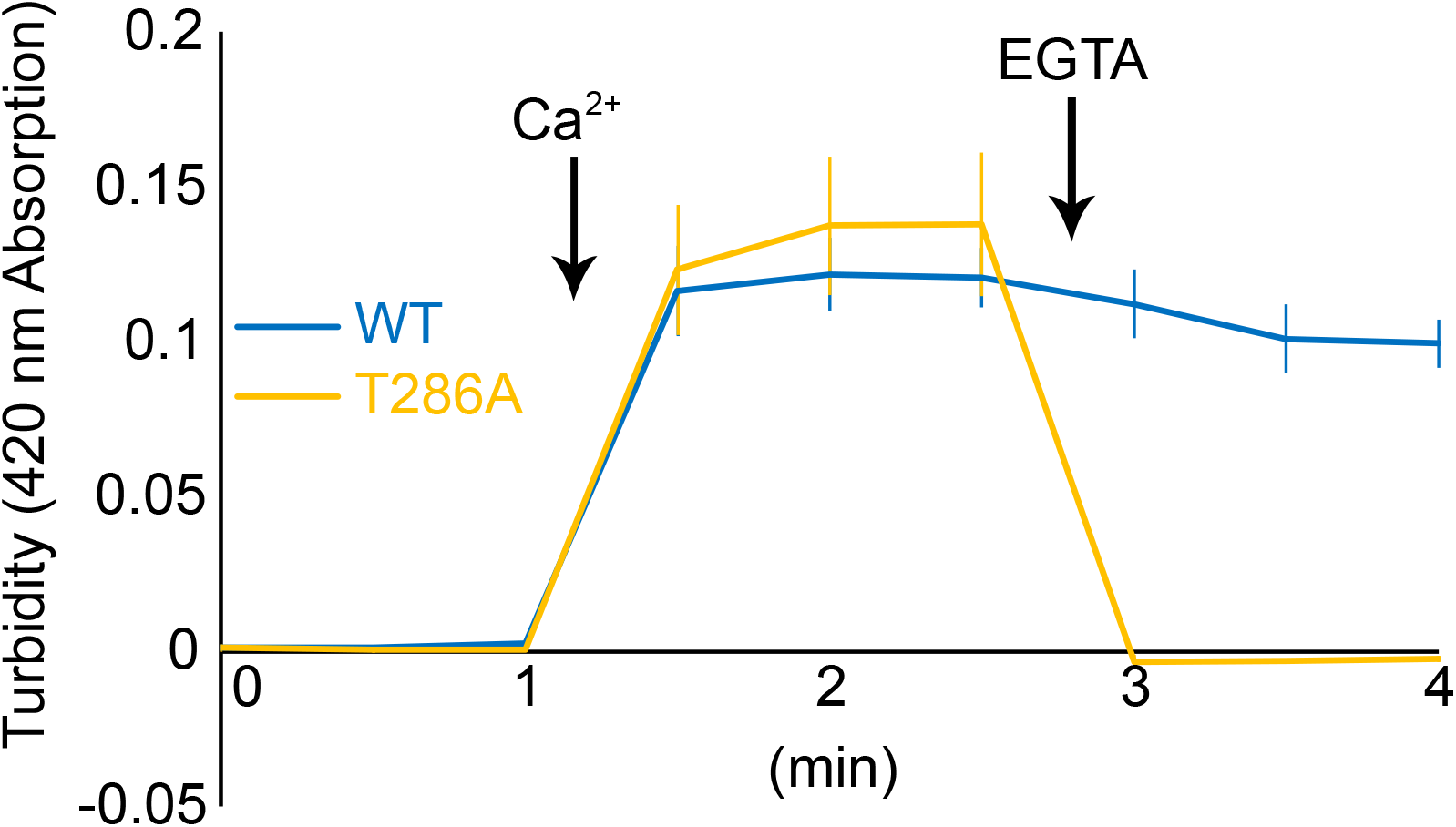
Turbidity measurement for CaMKII-GluN2Bc protein mixture. Protein mixture solution containing 10 μM CaMKII wildtype (WT) or T286A, 10 μM GluN2Bc and 10 μM calmodulin was subjected to turbidity measurement in the presence of ATP. 420 nm absorption before adding Ca^2+^ was defined as baseline. Turbidity was measured every 30 sec. CaCl_2_ and EGTA was added at indicated time point. N=4.

**Supplementary Figure 6.**
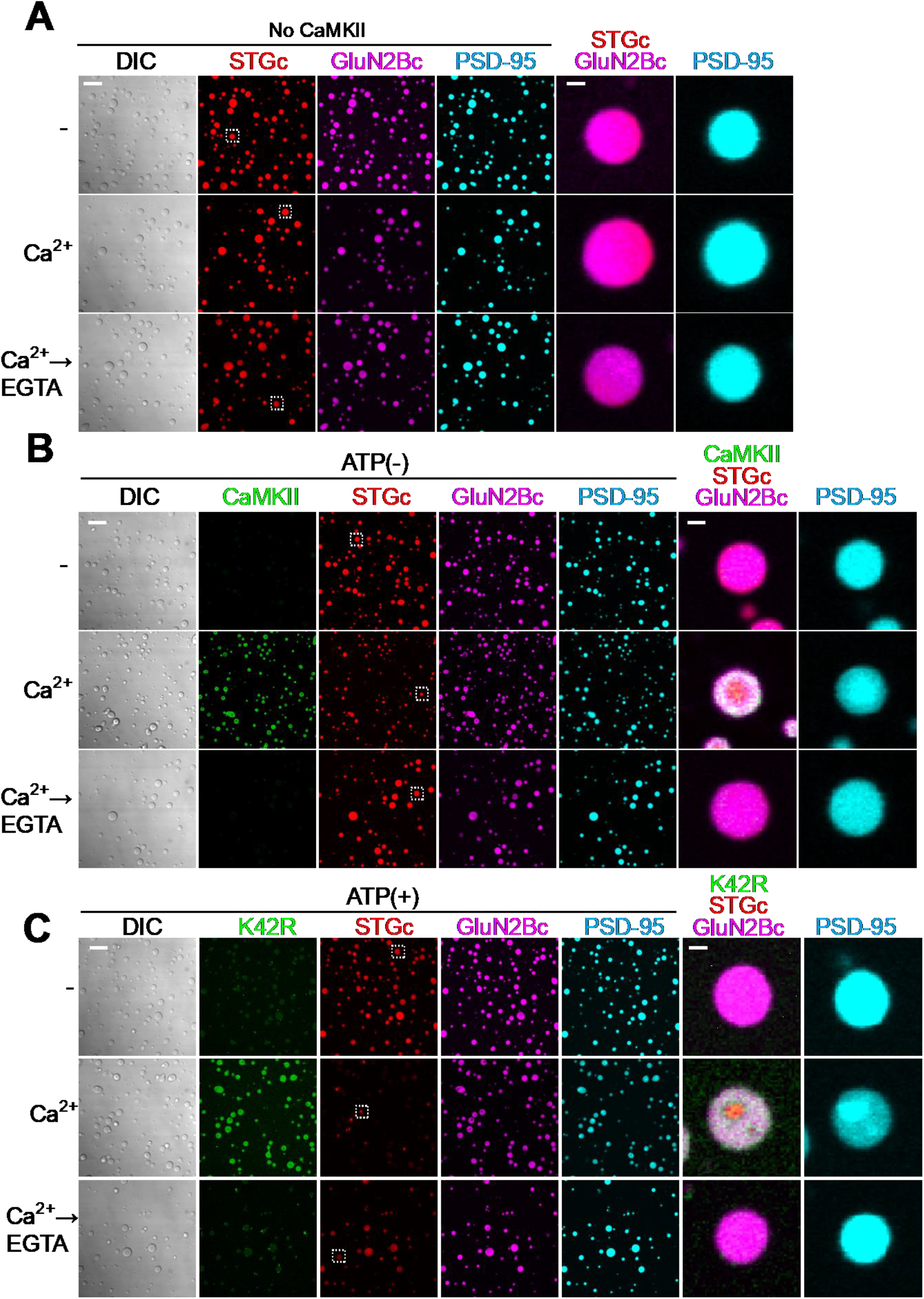
Persistent formation of the phase-in-phase assembly requires CaMKII and its kinase activity. (A) Images of the protein mixture consisting of 10 μM PSD-95, 7.5 μM STGc and 2.5 μM GluN2Bc in the presence of Mg^2+^-ATP but in the absence of CaMKII. This result indicates the requirement of CaMKII in the phase-in-phase organization formation. (B) Images of the protein mixture consisting of 10 μM PSD-95, 7.5 μM STGc, 2.5 μM GluN2Bc and 10 μM CaMKII in the absence of Mg^2+^-ATP. (C) Images of the protein mixture consisting of 10 μM PSD-95, 7.5 μM STGc, 2.5 μM GluN2Bc and 10 μM CaMKII K42R mutant in the presence of Mg^2+^-ATP. Right two columns are high magnification of dashed rectangle in STGc channel. Scale bars,10 μm and 1 μm for low- and high-magnification images. Phase-in-phase formation of STGc and PSD-95 condition was temporally observed in the presence of Ca^2+^ but upon chelation of Ca^2+^ with EGTA, the condensate returned to homogenous if ATP was removed (B) or catalytic activity of CaMKII is blocked by K42R mutation (C). Also, CaMKII was excluded from the phase. This indicates the requirement of catalytic activity of CaMKII for persistent formation of the phase-in-phase assembly in the absence of Ca^2+^.

**Supplementary Figure 7.**
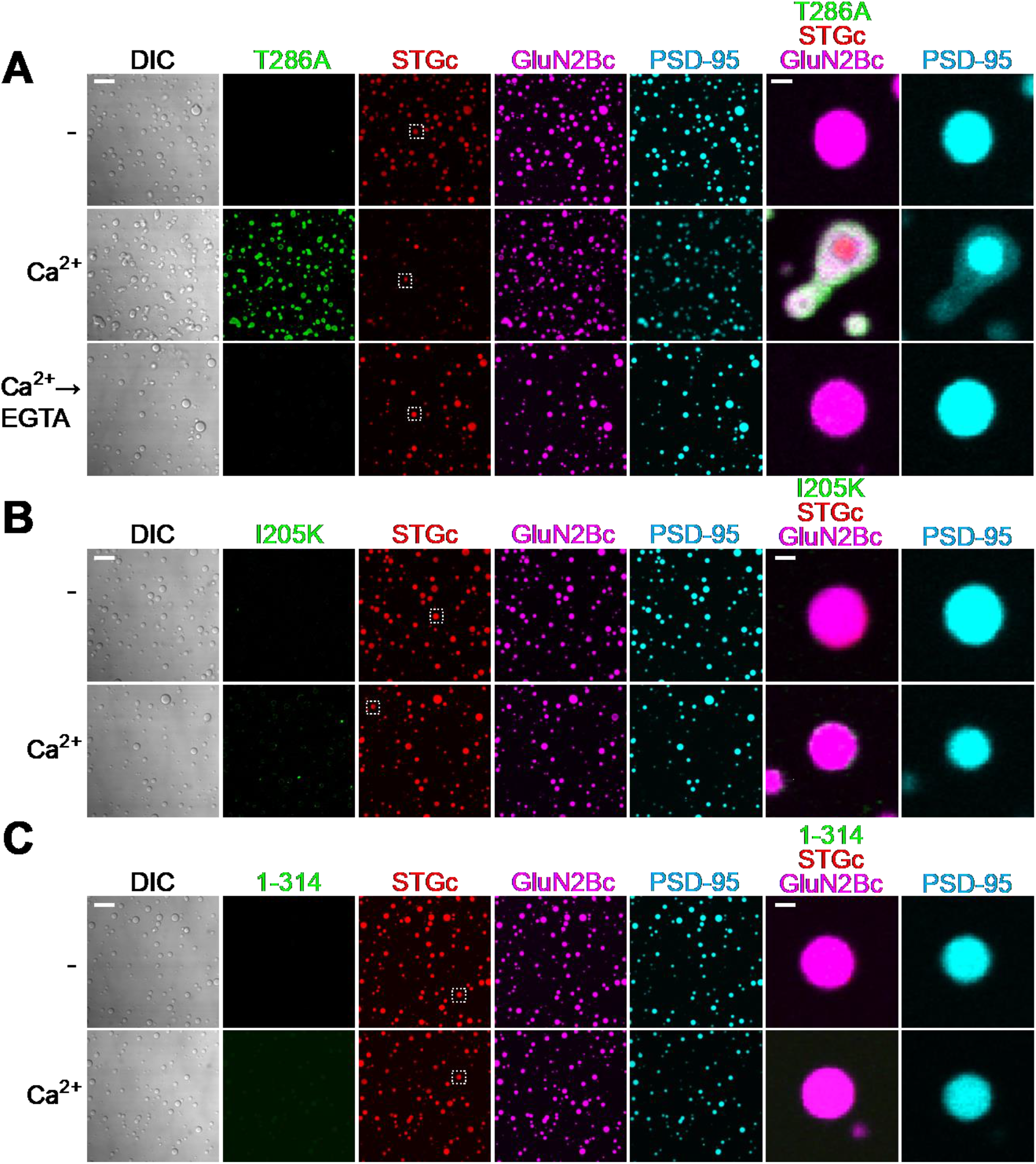
Persistent formation of the phase-in-phase assembly requires CaMKII autophosphorylation, T-site interaction, and multivalency. (A-C) Images of the protein mixture consisting of 10 μM PSD-95, 7.5 μM STGc, 2.5 μM GluN2Bc and indicated CaMKII mutant in the presence of Mg^2+^-ATP. -, Ca^2+^ and Ca^2+^→EGTA conditions are shown for T286A (A), and only - and Ca^2+^ conditions are shown for I205K (B) and 1-314 (C). High magnification images of the condensates are shown on the right. In (A), phase-in-phase formation of STGc and PSD-95 in CaMKII T286A condition was observed in the presence of Ca^2+^ but upon chelation of Ca^2+^ with EGTA, the condensate returned to homogenous. Also, CaMKII was excluded from the phase. This indicates the requirement of T286 phosphorylated CaMKII for persistent formation of phase-in-phase in the absence of Ca^2+^. (B) and (C) indicate the requirement of multivalent interaction between CaMKII T-site and GluN2Bc.

**Supplementary Figure 8.**
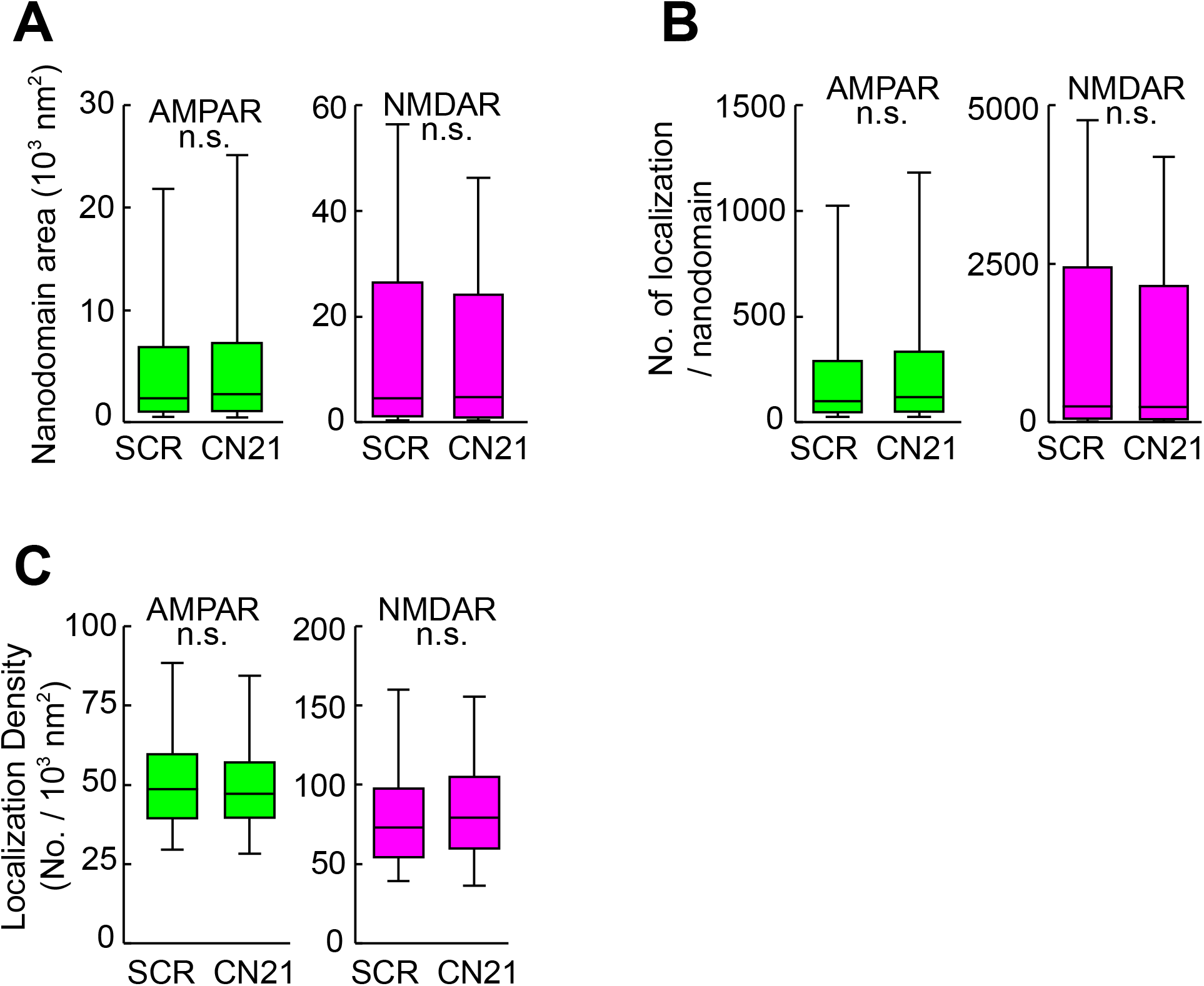
CN21 did not change the size and number of detected localization in the nanodomain. From dSTORM images, the area of each nanodomain (A, left, p=0.1677, right, p=0.4439), the number of detected localization per nanodomain (B, left, p=0.3826, right, p=0.4700) and the density of localization per area of nanodomain (C, left, p=0.1935, right, p=0.1069) were further analyzed using the same datasets as Figure 3C-E.

**Supplementary Figure 9.**
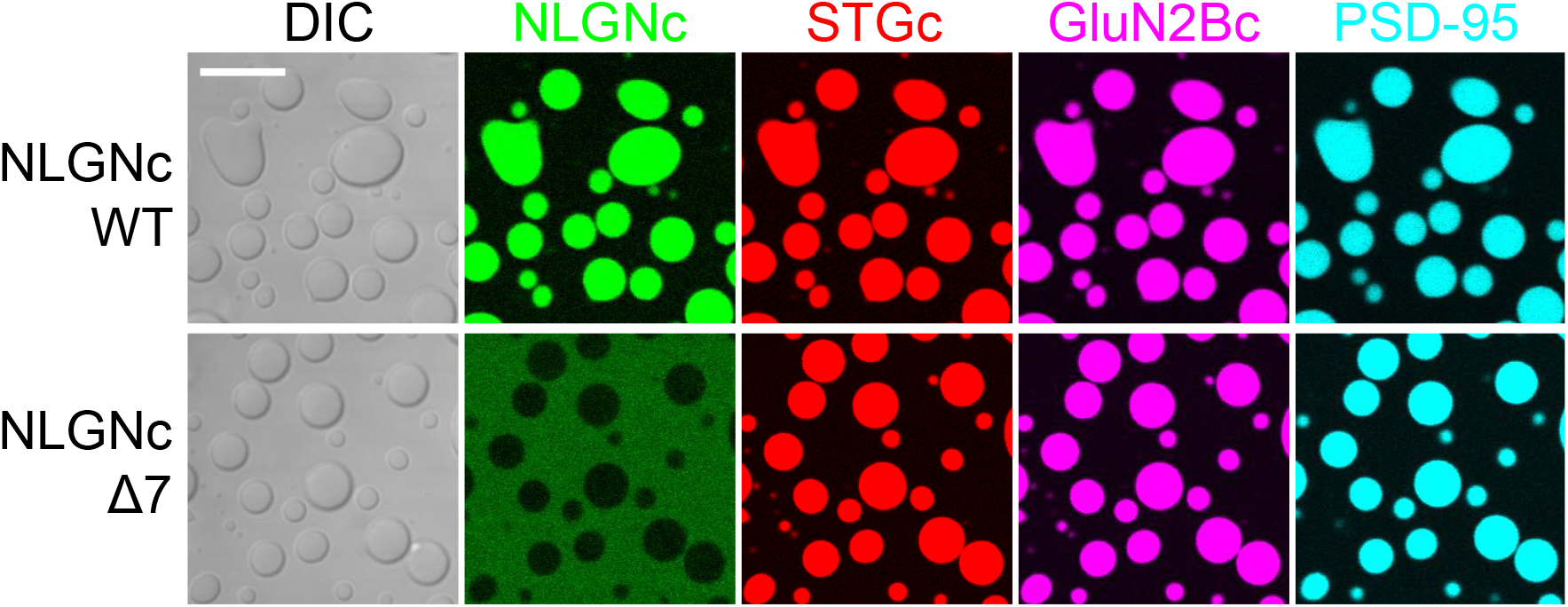
NLNGc participation in PSD-95 condensate requires PDZ-binding motif. Condensates were first formed by mixing purified PSD-95, GluN2Bc and STGc. Then either NLGNc wildtype (WT, top) or PBM deletion mutant (Δ7, bottom) were added. CaMKII was not added in here. Scale bar, 10 μm.

**Supplementary Figure 10.**
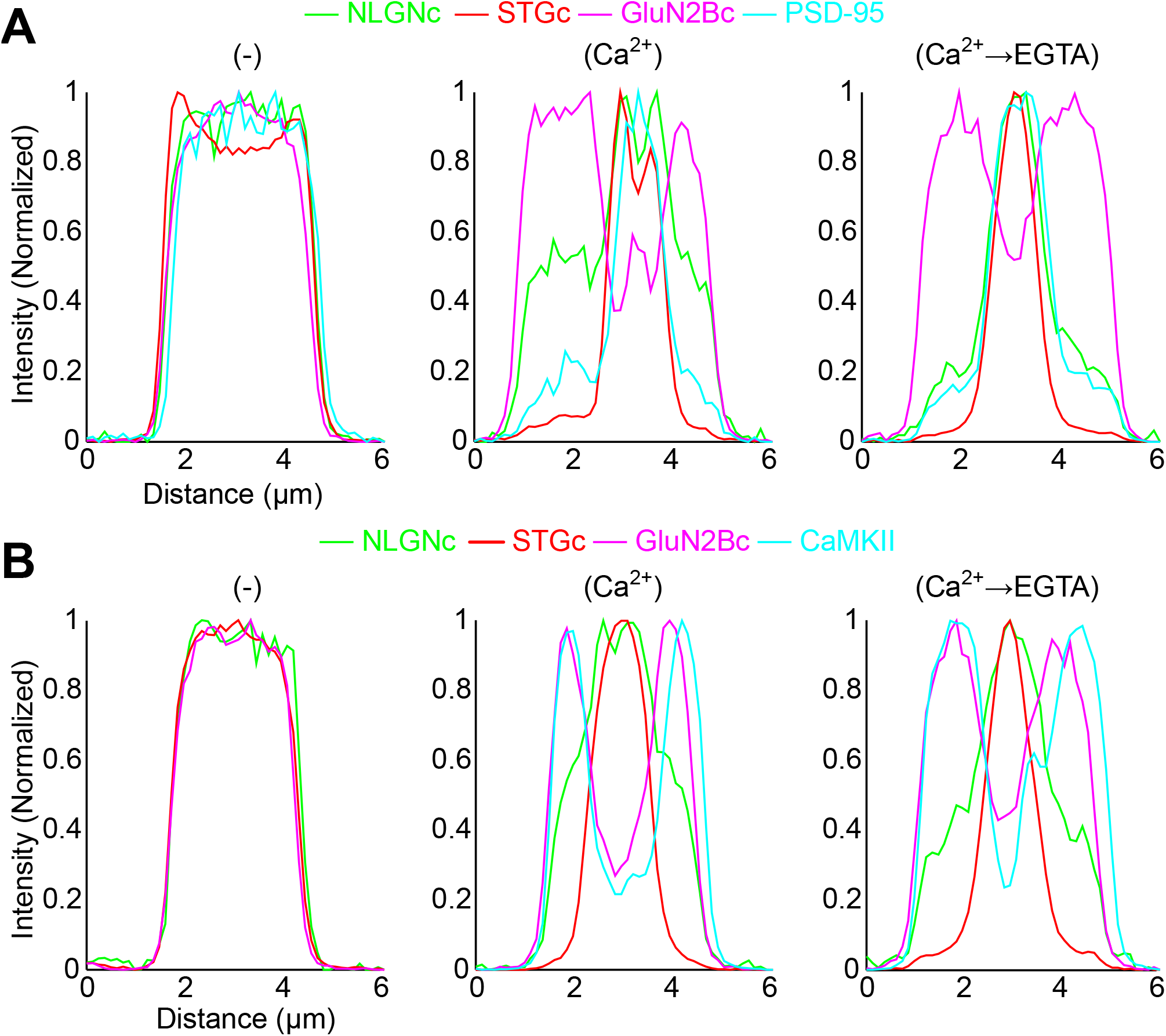
Fluorescence profiles of protein condensate in Figure 5. Graphs indicate the fluorescent profiles of the condensates formed by NLGNc (green), STGc (red), GluN2Bc (magenta) and PSD-95 (cyan, top) or CaMKII (cyan, bottom) in Ca^2+^ minus condition (left), Ca^2+^ condition (middle) and Ca^2+^→EGTA condition (right).

**Supplementary Figure 11.**
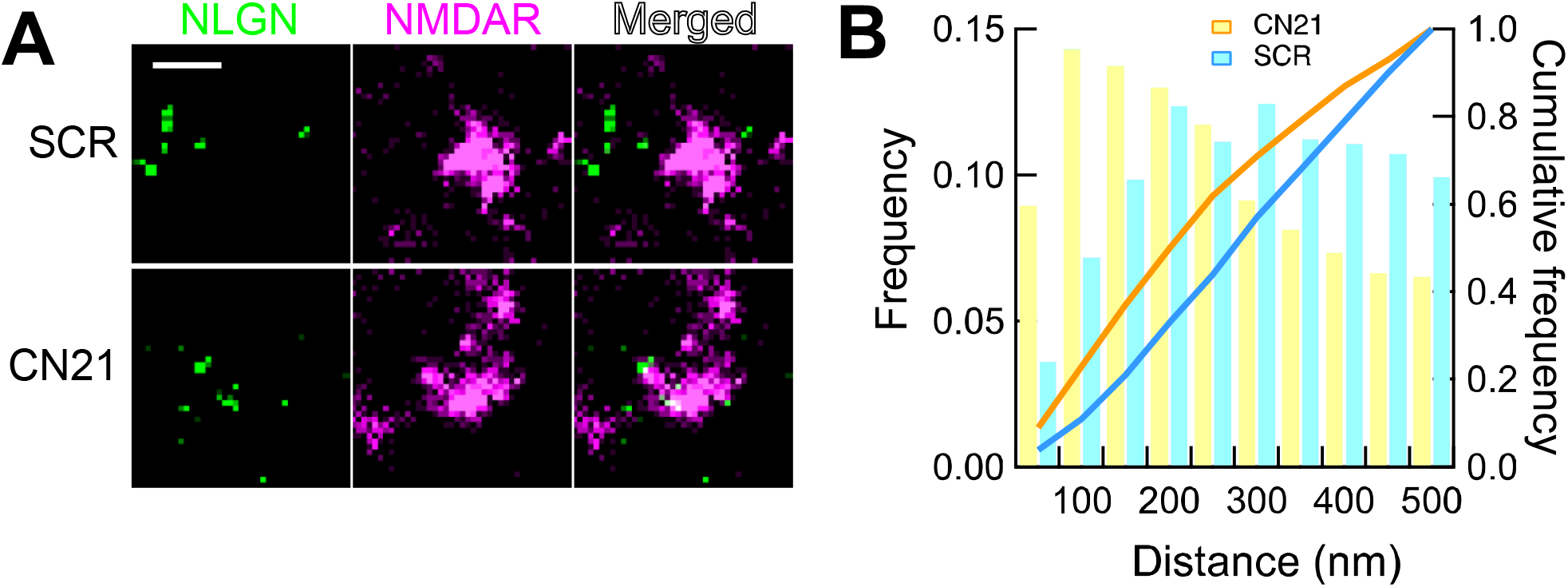
Effect of tat-CN21 on segregation between NMDAR and NLGN. A. Super-resolution (dSTORM) images of a synapse in cultured neuron, double-stained by NMDAR (GluN1 subunit) and NLGN. Hippocampal neurons were transfected with AP tag neuroligin and BirA. They were treated with 20 μM tat-scrambled (SCR) or tat-CN21 (CN21) for 30 min and labeled with monovalent streptavidin (to detect NLGN, green) and anti-GluN1 (NMDAR, magenta). Scale bar, 0.5 μm. B. The distribution of the distance from NMDAR localization to the nearest NLGN localization under two conditions. The frequency in each bin was normalized by the total number of localizations; CN21, 1733 and SCR, 1233. Statistical significance was tested by Kolmogorov-Smirnov test. α = 0.05; D = 0.180; critical value= 0.055; p = 6.83 ×10^-21^.

**Movie 1 Dispersion of CaMKII and GluN2Bc protein condensates by competing T-site interaction**

Time-lapse imaging of CaMKII-GluN2Bc condensates (Ca^2+^→EGTA condition) during infusion of 50 μM Camk2n1. Camk2n1 was manually infused from the top right of the image. At x 25 speed. See Fig. 3A for still images. Scale bars, 10 μm.

**Movie 2 Dispersion of CaMKII, GluN2Bc, PSD-95 and STGc protein condensates by competing T-site interaction**

Same experiment as in Movie 1 using the condensates with phase-in-phase formed by CaMKII, GluN2Bc, PSD-95 and STGc. PSD-95 was not imaged due to the limited number of color channels available. At x 50 speed. See Fig. 3B for still images. Scale bars, 10 μm.

## METHODS

### Guidelines

All recombinant DNA and animal experiments were carried out in accordance with the institutional guidelines of Kyoto University, Hong Kong University of Science and Technology, University of Bordeaux and CNRS.

### DNA constructs and protein purification

Rat CaMKII wild type and mutants, fluorescent proteins fused with Spy-catcher, Spy-tag fused with receptor C-tails such as GluN2Bc (mouse, a.a. 1226-1482), STGc (mouse, a.a. 203-323), NLGNc (mouse, a.a. 719-843), GluA1 (rat, a.a. 827 - 907), and GluA2 (rat, a.a. 834 - 883) were inserted into pSUMO vector. Amino acid residues were numbered with the initiation methionine as 1. PSD-95 and calmodulin were inserted into p32m3c vector as previously described ^13^.

All proteins were expressed in BL21 DE3 RIL strain and purified by affinity column using Nickel - NTA beads (Nacalai Tesque, Kyoto, Japan), gel filtration column HiLoad 26/600 Superdex 200 pg (GE healthcare, IL, USA) and anion exchange column HiTrap Q HP (GE Healthcare, IL, USA). All tags for purification were cut and removed. I205K mutant of CaMKII was tagged with GFP due to a difficulty of the expression and purification of untagged protein.

Fluorescent protein tagged Spy-catcher and Spy-tag tagged receptor C-tails were mixed with excess molar ratio of monomer C-tails and incubated for 2 hours at room temperature to covalently conjugate with each other. Extra monomer C-tails were removed by additional gel filtration. PSD-95 and CaMKII was labeled by iFluor 405 succinimidyl ester or iFluor 488 succinimidyl ester (AAT Bioquest, CA, USA) as previously described ^13^. Labeled protein was mixed with unlabeled protein at 1:100. Protein concentration is expressed as monomer concentration throughout the study.

### Formation and observation of LLPS condensates

Proteins were mixed in the buffer containing 50 mM Tris-HCl pH 7.5, 100 mM NaCl, 1 mM Tris(2-carboxyethyl) phosphine (TCEP), 0.5 mM EGTA and 10 μM calmodulin in the presence of 5 mM MgCl_2_ and 2.5 mM ATP (- condition). MgCl_2_ and ATP were not added in Fig. 1F and Fig. S6B. Two mM CaCl_2_ was added to activate CaMKII (Ca^2+^ condition) and 10 seconds later 2.5 mM EGTA was further added to chelate Ca^2+^ (Ca^2+^→EGTA condition) to mimic a transient Ca^2+^ signal.

Sedimentation assay was carried out as previously described ^12–14^. The protein sample in a low protein binding tube (WATSON, Tokyo, Japan) was centrifuged at 10,000 g for 1 min. Pellet and supernatant was denatured by SDS loading buffer and adjusted to the same volume. Five μL of samples were loaded onto SDS–PAGE and visualized by Coomassie brilliant blue.

For confocal microscope imaging, a sample chamber was made between a coverslip (12 mm round coverslip, MATSUNAMI, Osaka, Japan) and a slide glass (FRC-04, MATSUNAMI, Osaka, Japan) separated by double-sided adhesive paper tape as a spacer. Five μl of protein mixture was injected into this space and the condensates were allowed to settle down to the bottom of coverslip for 5 minutes. Observation was performed by a confocal microscopy system (FLUOVIEW FV1200, Olympus, Tokyo, Japan). Images of each colour channel were obtained with excitation wavelength and bandpass filters as follows; 405 nm for iFluor-405 tagged PSD-95 or CaMKII, 488 nm for iFluor-488 tagged CaMKII or Kusabira Green-tagged NLGNc, 546 nm for DsRed2-tagged STGc and 647 nm for eqFP670 tagged GluN2Bc and E2-Crimson tagged GluA1c and GluA2c. Tiam1 peptide (mouse, a.a. 1540-1560) was labelled with fluorescein by NHS-ester at the amino terminus.

### Turbidity assay

Ten μM CaMKII, 10 μM GluN2Bc were mixed in the buffer containing 50 mM Tris-HCl pH 7.5, 100 mM NaCl, 1 mM Tris(2-carboxyethyl) phosphine (TCEP), 0.5 mM EGTA and 10 μM calmodulin in the presence of 5 mM MgCl_2_ and 2.5 mM ATP. The turbidity of protein sample as the optical density at 420 nm was measured by nanodrop ND-1000 (Thermo Fischer, MA, USA). The baseline was defined as zero, and the turbidity was measured every 30 sec for 4 min. Two mM CaCl_2_ was added between 1 to 1.5 min and 2.5 mM EGTA was further added between 2.5 to 3 min.

### Cell culture, drug treatment and Immunostaining

Banker type cultures of hippocampal neuron were prepared from embryonic day 18 (E18) Sprague-Dawley rats at a density of 200,000 cells per dish as described ^4, 41^. The neurons at 16 days *in vitro* were treated with a CaMKII inhibitor peptide CN21 fused with a cell-penetrating peptide TAT (TAT-CN21; YGRKKRRQRRRKRPPKLGQIGRSKRVVIEDDR) or a scrambled CN21 with TAT (TAT-scrambled; YGRKKRRQRRRVKEPRIDGKPVRLRGQKSDRI) (20 μM) for 30 mins. After the treatment, the neurons were surface immunolabeled for endogenous glutamate receptor labeling: GluA2 (anti-GluA2, clone 14B11, 0.0033 μg/μl. IgG2b. From Dr. Eric Gouaux) and GluN1 (anti-GluN1, clone 10B11, 0.002 μg/μl. IgG1. From Dr. Gouaux) at 37 °C for 15 mins. Neuroligin-1 was labeled with biotin by co-expressing acceptor peptide (AP)-tagged neuroligin-1 and a biotin ligase BirA ^42^. The cells were surface labelled by incubating with monovalent streptavidin coupled to Alexa 647 for 10 min. After 3 washes, the cells were fixed with 4% paraformaldehyde (Sigma-Aldrich, #P6148) / 4% sucrose (Sigma-Aldrich, #0389) in phosphate buffered saline (PBS) at room temperature for 10 mins and treated with blocking solution (1.5% bovine serum albumin (Sigma-Aldrich, #A3059) / 0.1% fish skin gelatin / 0.1% Triton X-100 in PBS/NH4Cl) at room temperature for 1 hr. Cells were then incubated with secondary antibodies, goat anti-mouse IgG2b Alexa 647 (Thermo Scientific #21242) and goat anti-mouse IgG1 Alexa 532 (Thermo Scientific, and coupling done at IINS) at RT 1 hr. Following 3 washes, a second fixation was performed and then cells were imaged.

### Direct STochastic Optical Reconstruction Microscopy (dSTORM) imaging

dSTORM imaging was performed on LEICA DMi8 microscope equipped with Leica HCX PL APO 160x 1.43 NA oil immersion TIRF objective and fiber-coupled laser launch (532 nm and 642 nm) (Roper Scientific, Evry, France). Single fluorophores were detected with EMCCD camera (Evolve, Photometrics, Tucson, USA). Sample was mounted on a Ludin chamber (Life Imaging Services, Switzerland) and 600 μl dSTORM pyranose switching buffer ^43^ was added. An additional coverslip was placed on top to minimize buffer evaporation and oxygen exchanges with ambient air. Before dSTORM imaging, a diffraction limited image of the target region (512 x 512 pixels, 1 pixel=100 nm) was taken under wide-field epifluorescence illumination. Image acquisition was steered by MetaMorph software (Molecular Devices, USA) with 30 ms exposure time, 20,000 frames per each color. The 642 nm and 532 nm lasers were used sequentially. Multi-color fluorescent microspheres (Tetraspeck, Invitrogen, #T7279) were used as fiducial markers for nanometer scale lateral drifts correction and dual color registration.

### Nanodomain analyses

To analyze AMPAR and NMDAR nanodomains, intensity super-resolution images with a pixel size of 25 nm were reconstructed during the acquisition using WaveTracer software operating as a plugin of MetaMorph ^44^. Lateral drifts were corrected automatically from the localizations of fluorescent fiduciary markers absorbed into the coverslip. Single molecule localizations of Alexa 532 and Alexa 647 were aligned post-acquisition with PALMTracer software using a 3^rd^ order polynomial transform to correct for chromatic aberrations on the whole field of view. SR-Tesseler and Coloc-Tesseler tessellation-based analysis software ^45, 46^ were used to quantify respectively the nanodomains and the colocalization of AMPAR and NMDAR. The segmentation of AMPAR and NMDAR nanodomains was performed separately. Single molecule localizations were used to compute the Voronoï tessellation, from which the 1^st^ rank order local density map was computed. Clusters were segmented automatically using a threshold of twice the average local density of the whole dataset, with a minimum localizations number of 5 and a minimum area of 2 × 10^4^ nm^2^.

Next, the clusters’ nanodomains were segmented by applying a threshold of one time the average density within each cluster, with a minimum localizations number of 25, a minimum area 0.02 (AMPAR) or 0.01 (NMDAR) × 10^4^ nm^2^. The colocalization between AMPAR and NMDAR was computed from the overlapping nanodomains area within selected regions of interest (ROI). ROIs were identified from merged epifluorescence images of AMPAR and NMDAR.

NLGN1 and NMDAR double stained images were analyzed similarly but because they hardly overlapped, we measured the distance from NMDAR localization to the nearest NLGN1 localization with a cut-off of 500 nm. Statistical significance was tested by Kolmogorov-Smirnov test. α was set at 0.05.

## Acknowledgement

We thank Drs. Roger A. Nicoll, Johannes W. Hell, and Thomas A. Blanpied for comments on the manuscript, Drs. Eric Gouaux, Olivier Thoumine, and Matthieu Sainlos and Bordeaux Imaging Center for the reagents and Dr. Lily Yu, Adam Z. Weitemier, and Ms. Emily Agnello for editing.

## Author contributions

T. H. and P. L. conducted and managed all experiments. Y. H. managed the overall project. Q. C. and M. Z. participated in LLPS experiments. J. F., F. L., C. B., J. S., D. C., L. G. and E. H. participated in super-resolution microscopy.

